# Identification, Purification and Characterization of Mast Cells in Murine Liver Fibrosis: Novel Methods, Expression Signatures and Correlation with Severity

**DOI:** 10.1101/2025.07.25.666577

**Authors:** Christian Penners, Julia Otto, Steffen K. Meurer, Ralf Weiskirchen, Michael Huber, Christian Liedtke

## Abstract

Mast cells (MCs) are myeloid cells of the innate immune system. As a first line of defense, they fulfill effector functions and immune modulatory properties. Upon activation, they release pro-inflammatory mediators such as cytokines and proteases. It has been suggested that MCs may contribute to the development of liver fibrosis. However, investigating hepatic MC biology in mice is challenging due to low MC numbers and a lack of suitable detection techniques relying on MC proteins and their modifications. Here, we evaluated whether the expression strength of MC markers correlates with the degree of liver fibrosis in mice and aimed to determine the frequency and localization of hepatic MCs. We applied both a toxic (DEN/CCl_4_ treatment) and a genetic (Mdr2^-/-^ mice) liver fibrosis model in C57BL/6 mice and found significant correlations between fibrosis grade and the expression of several established MC markers. These correlations were further supported by data analyses from patients with fibrosis and hepatocellular carcinoma (HCC) using publicly available transcriptomics datasets. We used FACS to purify and isolate MCs from fibrotic mouse livers and verified MC signatures by qPCR analysis of MC-specific gene expression. Hepatic MCs were predominantly negative for Mast Cell Protease 5 (*Mcpt5*) and occurred at a low frequency (approximately 1-2% of leukocytes). Using Molecular Cartography^TM^ of fibrotic liver sections, we determined the spatial localization, expression signature, abundance (approximately 2 cells/mm^2^) and cellular environment of murine hepatic MCs.

In summary, we demonstrated the existence of MCs in murine fibrotic livers and defined MC expression signatures that correlate with the strength of liver fibrosis. These findings will help to study MC biology in murine models of liver disease more effectively in the future.

## 1 Introduction

Mast cells (MCs) are myeloid cells best known for their essential roles in allergic inflammation (1). They are typically found in barrier tissues exposed to the environment such as the skin, lungs, and gastrointestinal tract, but have also been described in other vascularized organs such as the kidney and liver (2). MCs are activated by antigen/allergen crosslinking of immunoglobulin E (IgE)-loaded high-affinity IgE receptors (FcεRI). Upon activation, MCs release a wide range of preformed inflammatory mediators, including histamine and proteases, as well as de-novo generated mediators such as arachidonic acid metabolites and cytokines (3). The development and survival of MCs depend on the stem cell factor (SCF)/KIT signaling axis. KIT, a receptor tyrosine kinase, is expressed at high levels on MCs and is commonly used as a lineage marker along with FcεRIα (encoded by *Fcer1a*) (2). MCs are subclassified as mucosal MCs (MMCs) and connective tissue-type MCs (CTMCs), distinguished by anatomical location, cytokine dependency, and granule protease expression (4, 5). MMCs characteristically express *Mcpt1* and *Mcpt2*, and develop under the influence of IL-3 and TGF-β (6–8), whereas CTMCs preferentially express *Mcpt4, Mcpt5* (also known as *Cma1*), *Tpsb2* (tryptase β2), (8) and *Cpa3* (carboxypeptidase A3) (9). Recent studies in humans suggest an involvement of MCs in hepatic fibrogenesis. For example, human liver biopsies from patients with advanced primary sclerosing cholangitis (PSC) exhibited marked MC infiltration, detectable by immunohistochemistry using toluidine blue staining (10). However, the existence and precise role of MCs in murine models of liver fibrosis remain unclear due to the low abundance of resident MCs, limited availability of mouse-specific MC antibodies and contradictory data. In this study, we aimed to convincingly demonstrate the existence and abundance of MCs in the fibrotic liver and correlate them with fibrosis progression. Moreover, our aim was to analyze if MCs play a role in disease progression. To achieve this, we utilized spatial transcriptomics analysis to detect MCs in sections of fibrotic mouse livers. We conducted correlation analyses between MC markers and fibrosis progression in three different murine fibrosis models and compared these findings with data from patients. Our results demonstrate a significant correlation between the expression of MC markers and the severity of fibrosis progression in mice. Additionally, we outline a protocol for isolating primary liver MCs from mice.

## 2 Material and methods

### 2.1 Maintenance of mice and general animal experimentation

Animal experiments were conducted in compliance with German legal requirements and animal protection laws, approved by the authority for environment conservation and consumer protection of the state of North Rhine-Westphalia (LANUV, Recklinghausen). Mice of both genders were housed in a temperature-controlled room with 12-hour light/dark cycles and had free access to food and water. All mice used in this study were maintained in a C57BL/6 background.

For correlation analyses between the degree of liver fibrosis and MC markers, we aimed to include as many samples as possible from different fibrosis models with a high degree of variability in the severity of fibrosis. In accordance with the 3R principles for minimizing animal testing (11), we also included fibrotic liver samples from previous experiments. We used two established mouse models of experimental liver fibrosis: In the toxin-induced DEN/CCl_4_ model (12), wildtype (WT) mice were treated with a single injection of *N*-nitrosodiethylamine (DEN, 25 mg/kg, *i.p.*) in 14-day-old pups followed by weekly injections of carbon tetrachloride (CCl_4_, 0.5 ml/kg in corn oil, *i.p.*) starting at the age of 6 weeks for 18 weeks. For these analyses, some fibrotic samples from DEN/CCl_4_-treated mice lacking the *Ccne1* gene exclusively in *Lrat*-expressing cells were included as documented in Supplementary Table 1.

As a second liver fibrosis model we applied genetic inactivation of the multidrug resistance protein 2 (Mdr2^-/-^) resembling cholestatic liver fibrosis (13, 14). To this end, we used liver samples from Mdr2^-/-^ mice at the age of 26 and 52 weeks, respectively. For these analyses, some fibrotic liver samples from Mdr2^-/-^ mice with additional deletion of the caspase-8 (*Casp8*) gene in hepatocytes (Mdr2^-/-^*Casp8*^Δhepa^) were included. In an earlier study, we demonstrated that depletion of *Casp8* has no effect on liver fibrosis progression in male Mdr2^-/-^ mice (15).

Double-fluorescent *Cre*-reporter mice (mT/mG) were purchased from The Jackson Laboratory (*B6.129(Cg)-Gt(ROSA)26Sor^tm4(ACTB-tdTomato,-EGFP)Luo^/J*, JAX stock Strain #:007676, The Jackson Laboratory, Bar Harbor, ME, USA). For this study, we generated mT/mG; *Mcpt5*-*Cre* mice (16, 17) and then crossed these mice with Mdr2^-/-^ mice (18) in a C57BL/6 background to generate Mdr2^-/-^ mT/mG; *Mcpt5*-*Cre* triple-transgenic animals. Cre-negative, floxed Mdr2^-/-^mT/mG littermates and C57BL/6 WT mice were used as controls.

### 2.2 Immunohistochemistry and staining of MCs in murine tissue

Explanted mouse livers and control organs were fixed in 4% paraformaldehyde (PFA) immediately after extraction, embedded in paraffin, sectioned and subjected to staining for Hematoxylin/Eosin (H&E), Alcian blue or Toluidine blue as previously described (6).

### 2.3 RNA isolation and quantitative Real-Time PCR (qPCR)

Total RNA from liver tissues or cell pellets was isolated using the QIAzol Lysis Reagent (Qiagen, Hilden, Germany). Reverse-transcription was performed using an Omniscript RT Kit (Qiagen, Hilden, Germany). Relative quantitative gene expression was measured *via* real-time PCR using a QuantStudio 5 and 6 Flex Real Time PCR System (Applied Biosystems, Foster City, CA, USA) and a Fast SYBR Green PCR master mix (Invitrogen, Carlsbad, CA, USA). Target gene expression was normalized to *Gapdh* expression as internal standard and calculated as fold induction compared to controls as indicated in the figure legends. Primer sequences used in this study are listed in Table 1.

**Table 1.**
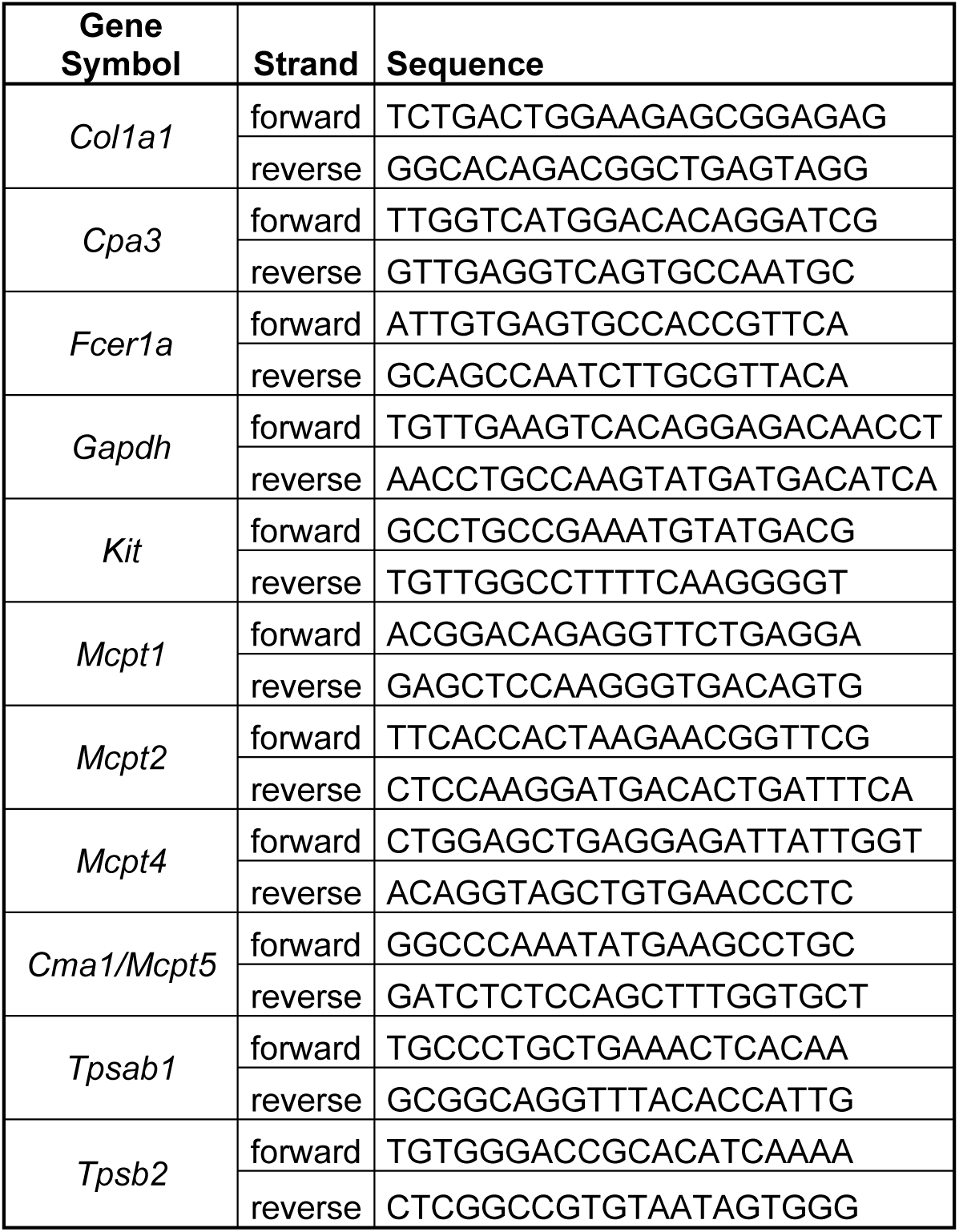
List of used primer sequences.

### 2.4 Isolation and flow cytometry of MCs

Bone marrow derived mast cells (BMMCs) were generated from immature MC precursors of the bone marrow as recently described (19). Hepatic MCs from mice were isolated as follows: Hepatic immune cell suspensions were obtained by digesting whole minced livers for 1 h at 37 °C in DMEM containing 1.6 mg/ml collagenase IV (Worthington, Lakewood, USA) and 0.4 mg/ml DNAse I (Roche, Basel, Switzerland). The cell suspension was filtered through a 70 µm cell strainer, hepatocytes were removed by low-speed centrifugation and red blood cells were lysed using RBC lysis buffer (ThermoFisher, Waltham, USA). Cells were stained for flow cytometry using 1:100 dilutions of CD45 - APC-Cy7, CD11b - V450, CD117 – PE-Cy7 (all BD, Franklin Lakes, USA), CD11c - eFluor 506, FCER1A - APC (both ThermoFisher, Waltham, MA, USA) followed by either flow cytometry using a BD LSR Fortessa or FACS sorting using a BD FACSAria Fusion.

### 2.5 Bioinformatic analysis

Human mRNA sequencing data from tumorous or normal liver tissue were obtained from the TCGA-LIHC dataset of the TCGA Research Network: https://www.cancer.gov/tcga. Murine mRNA sequencing data from fibrotic livers induced by chronic administration of CCl_4_ were obtained from publicly available data on GEO (GSE167216) (20). Analysis was performed using R v4.4.2. Normalized counts were obtained using variance stabilizing transformation (VST) from the DESeq2 package v1.46.0 (21), followed by analysis of the Pearson correlation coefficient and significance of correlation using the built in cor.test function. Correlation matrices were generated using the GGally package v2.2.1. Differential gene expression was analyzed between tumor samples from the TCGA-LIHC dataset with high or low expression of a mast cell gene signature. VST normalized counts were utilized to generate an expression score for each sample based on the mast cell specific genes *Cpa3*, *Tpsab1* and *Tpsb2* using the GSVA package v2.0.7 (22). Samples were classified into low or high expression of the mast cell signature based on the median score followed by differential gene expression analysis using DESeq2. Gene ontology enrichment on significant upregulated genes was analyzed using the clusterProfiler package v4.14.6, (23) and visualized using the enrichplot package v1.26.6.

### 2.6 Molecular Cartography™

Molecular Cartography™ was conducted at Resolve BioSciences GmbH (Monheim, Germany), following the protocol outlined by Guilliams et al. (24) with minor adjustments. Formalin-fixed paraffin-embedded (FFPE) tissue sections from Mdr2^-/-^ mice at 26 and 52 weeks of age, exhibiting advanced liver fibrosis, were used for the analysis. Sections were deparaffinized before the Molecular Cartography procedure. Probe design for mast cell-specific genes was conducted according to Guilliams et al. (24). The complete list of target genes, along with the respective catalogue numbers provided by Resolve Biosciences, can be found in Table 2.

**Table 2.** Gene panel for Molecular Cartography^TM^.

| Gene | Resolve Catalog Number |
| --- | --- |
| *Cd200r3* | Q0VAH |
| *Cma1* | P07H9 |
| *Cpa3* | PZR6E |
| *Fcer1a* | QQT A7 |
| *Il1rl1* | P6E70 |
| *Kit* | PQE4E |
| *Mcpt1* | Q9VAT |
| *Mcpt2* | Q2VAK |
| *Mcpt4* | Q1VAJ |
| *Mrgprb2* | QGPA4 |
| *Tpsab1* | QWTAD |
| *Tpsb2* | Q7PAX |

The analysis of Molecular Cartography™ data was carried out as follows: DAPI images, transcript coordinates and cell segmentation data provided by Resolve Biosciences were processed using ImageJ2 and the Polylux v1.9.0 plugin to create overlays of cells and individual transcripts. Mast cells were identified as cells expressing *Kit* or *Cpa3*, along with at least one additional MC marker gene (*Cd200r3, Cma1, Fcer1a, Il1rl1, Mcpt1, Mcpt2, Mcpt4, Mrgprb2, Tpsab1,* or *Tpsb2*). The total number of MCs per section was then quantified.

### 2.7 Statistical analysis

Statistical significance of the results was tested using GraphPad Prism 10 (GraphPad Software, Inc., USA) for qPCR and FACS data, and R with the built-in cor.test function and ggpubr package (v0.6.0) for the analysis of transcriptomics data. Comparisons between two groups were performed using either an unpaired *t*-test in the case of normally distributed results or a Mann-Whitney test in the case of non-normally distributed results. Correlations between two variables were calculated by Pearson correlation together with a two-tailed *p*-value. Data represent mean ± standard deviation (SD) unless otherwise stated. In some cases, data are displayed as boxplots with the central line indicating the median, box edges representing the interquartile range (IQR), and whiskers extended to 1.5-fold of IQR. Individual points beyond the whiskers are considered outliers. In general, significance was defined as *: *p* ≤ 0.05; **: *p* ≤ 0.01; ***: *p* ≤ 0.001; ****: *p* ≤ 0.0001.

## 3 Results

### 3.1 MCs in the mouse liver cannot be identified using established histologic staining techniques

In a previous study, MCs were detected in human biopsies from patients with advanced primary sclerosing cholangitis (PSC) using immunohistochemical methods such as toluidine blue staining (6). In the present study, the goal was to better understand MC biology in the liver by using mouse models of liver fibrosis. Initially, the aim was to stain murine hepatic MCs in fibrotic/tumorous livers of Mdr2^-/-^ mice with alcian blue or toluidine blue, as done in human liver studies (6).

Using this method, murine MCs were found in the hilum and skin of the ear. However, they could not be detected in the liver parenchyma of Mdr2^-/-^ mice with pronounced liver fibrosis (Supplementary Figure 1A-C). Concomitant expression of KIT and FCER1A is a property of human and murine MCs (2). However, cells co-expressing KIT and FCER1A could not be detected in the mouse liver using immunofluorescence (data not shown).

### 3.2 Expression of MC-specific genes correlates with the degree of liver fibrosis in mice

In the present study, we tested the hypothesis that MCs in the mouse liver are associated with the severity of liver fibrosis. Since direct detection of MCs in the mouse liver by conventional staining techniques did not work in our hands, correlation studies were performed to obtain initial evidence for the involvement of MCs in liver fibrogenesis. For this purpose, we recruited mouse cohorts from several liver fibrosis models and their respective healthy controls (untreated or wildtype), providing a broad spectrum of liver samples with a high variability in fibrosis severity.

We treated mice with diethylnitrosamine (DEN) and carbon tetrachloride (CCl_4_) to generate liver samples with different severity of fibrosis and HCC (Figure 1A). The severity of liver fibrosis was quantified by determination of *Col1a1* expression showing clear induction by DEN/CCl_4_ with high variability in comparison to untreated controls as anticipated (Figure 1B).

**Figure 1.**
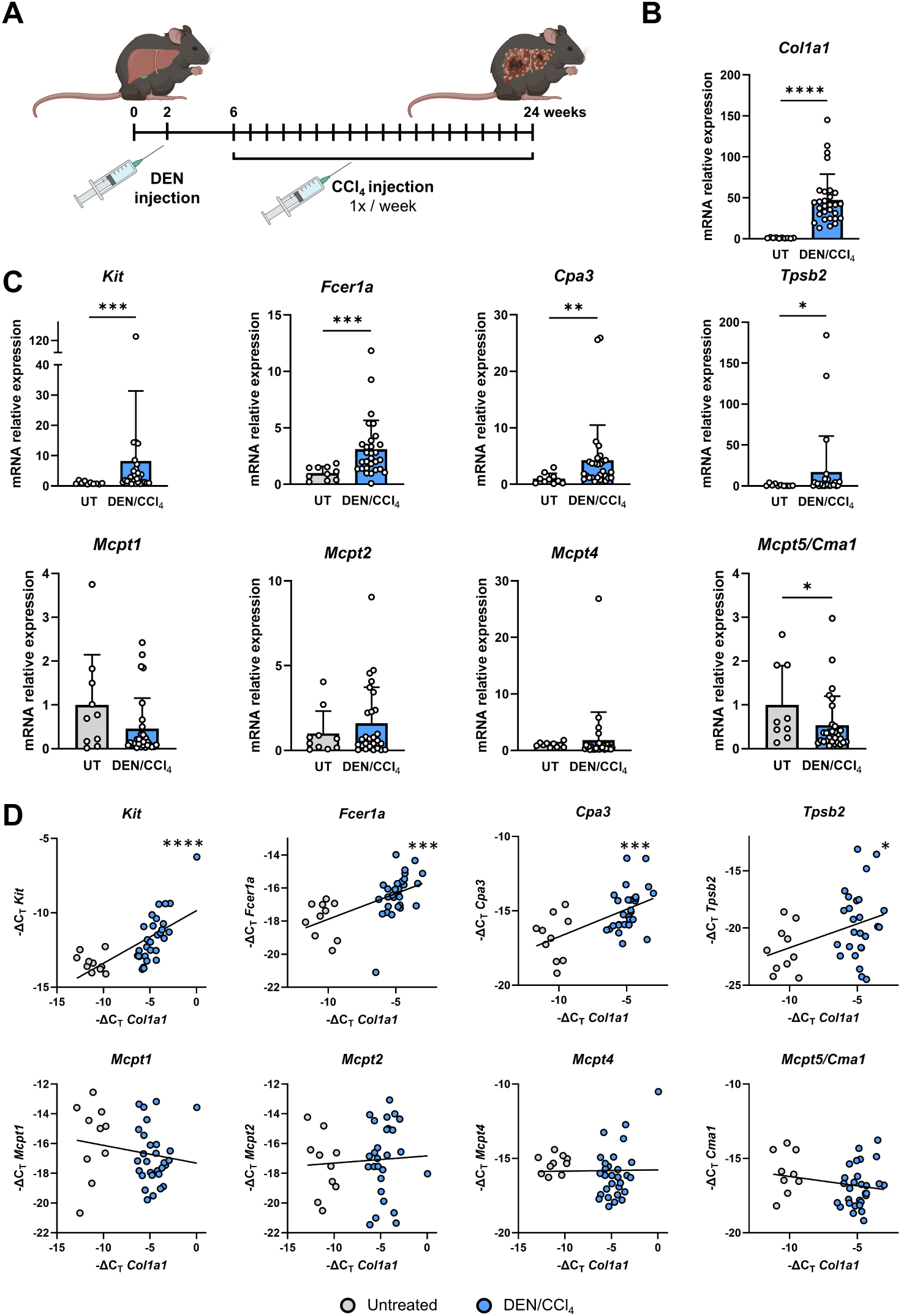
The severity of DEN/CCl_4_-induced liver fibrosis in mice correlates with gene expression levels of several MC markers. (**A**) Experimental setup: Mice were exposed to the DEN/CCl_4_ fibrosis/HCC model. Animals received a single injection of DEN on day 14 after birth to induce HCC, followed by weekly CCl_4_ administration starting at 6 weeks of age to induce liver fibrosis. Animals were euthanized at 24 weeks of age for fibrosis analysis. Untreated mice (UT) were used as controls. (**B**) Relative gene expression of *Col1a1* in untreated and DEN/CCl_4_ - treated mouse livers determined by qPCR. Gene expression was normalized to *Gapdh* expression and calculated as fold induction compared to untreated mice. (**C**) mRNA expression of MC-specific genes normalized to *Gapdh* expression and calculated as fold induction compared to untreated controls. Data presented as mean ± SD. (**D**) Pearson correlation analysis of *Col1a1* expression (as a measure of liver fibrosis) with the expression of MC-specific genes (*Kit*, *Fcer1a*, *Cpa3*, *Tpsb2*, *Mcpt1*, *Mcpt2*, *Mcpt4*, *Mcpt5*) determined by qPCR. Gene expression normalized to *Gapdh* and shown as -ΔC_T_. Samples with undetermined C_T_-values were excluded from the analysis; colour code indicates treatment regimen of each data point. *: *p* ≤ 0.05, **: *p* ≤ 0.01, ***: *p* ≤ 0.001, ****: *p* ≤ 0.0001.

Next, we used these liver samples to investigate the expression of genes that are typically expressed in MCs, such as *Kit*, *Fcer1a*, Carboxypeptidase A3 (*Cpa3*), Tryptase beta 2 (*Tpsb2*) and Mast cell proteases (*Mcpt*) 1, 2, 4 and 5 (*Mcpt5* is also referred to as Chymase 1 or *Cma1*). Our results demonstrated that the MC markers *Kit*, *Fcer1a*, *Cpa3* and *Tpsb2* were significantly up-regulated in liver samples obtained from mice treated with DEN/CCl_4_. Furthermore, the data indicated that *Mcpt5*/*Cma1* was generally down-regulated in fibrotic livers, while *Mcpt1, Mcpt2* and *Mcpt4* did not show aberrant expression in liver fibrosis (Figure 1C). Pearson correlation analysis using untreated and DEN/CCl_4_-treated liver samples revealed a significant correlation between *Col1a1* expression and the expression of *Kit*, *Fcer1a*, *Cpa3* and *Tpsb2* (Figure 1D). Importantly, the expression of *Mcpt1*, *Mcpt2*, *Mcpt4* and *Mcpt5*/*Cma1* did not correlate with the severity of fibrosis (Figure 1D). Altogether, this data suggests that MCs with a specific gene expression signature are present in the mouse liver and increase in number during fibrosis progression.

We wanted to confirm these findings in a second independent mouse model of cholestatic liver fibrosis. To do this, we utilized the Mdr2^-/-^ mouse model (25). Mice with a deletion of the Mdr2/*Abcb4* gene, which encodes multidrug resistance protein 2, spontaneously develop cholestasis and severe cholestatic liver fibrosis that worsens with age (Figure 2A).

**Figure 2.**
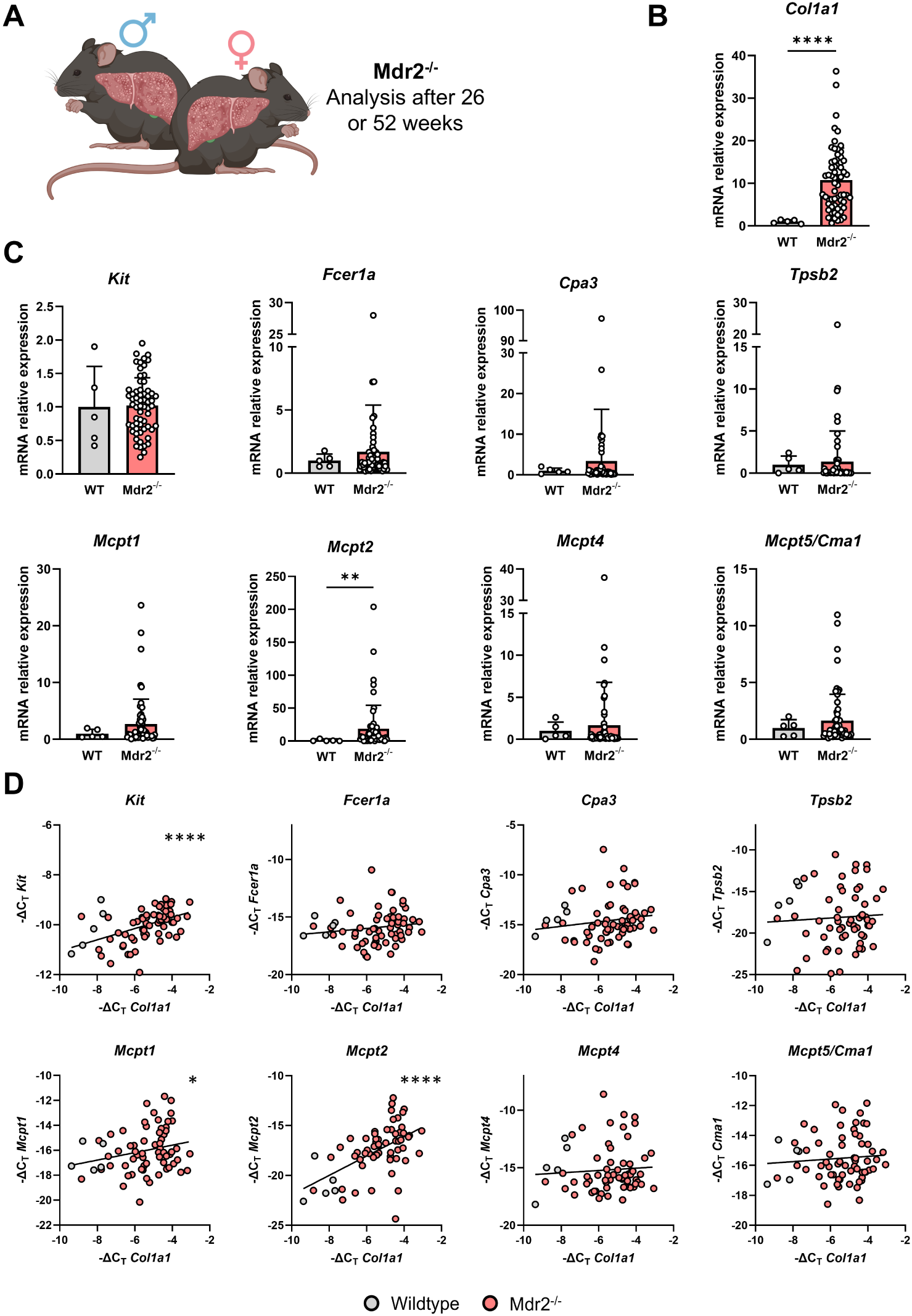
Gene expression levels of MC markers correlate with the severity of spontaneous cholestatic liver fibrosis in Mdr2^-/-^ mice. (**A**) Mice lacking the *Abcb4* gene (referred to as Mdr2^-/-^) develop liver fibrosis during ageing. To represent a wide range of fibrotic Mdr2^-/-^ livers, we included male and female Mdr2^-/-^ and WT mice aged 26 and 52 weeks in this analysis. Expression data and genotypes for each experimental mouse are listed in Supplementary Table 1. (**B**) The induction and severity of liver fibrosis were assessed by analysing the relative gene expression of *Col1a1* in WT and Mdr2^-/-^ livers at 26 and 52 weeks of age by qPCR. Gene expression was normalized to *Gapdh* expression and presented as fold induction compared to WT mice. (**C**) mRNA expression of MC-specific genes was determined, normalized to *Gapdh* expression and presented as fold induction compared to WT mice. Data are shown as mean ± SD. (**D**) Pearson correlation analysis of *Col1a1* expression (serving as a measure of liver fibrosis) with the expression of MC-specific genes (*Kit*, *Fcer1a*, *Cpa3*, *Tpsb2*, *Mcpt1*, *Mcpt2*, *Mcpt4*, *Mcpt5*) as determined by qPCR. Gene expression was normalized to *Gapdh* expression and presented as -ΔC_T_. Samples with undetermined C_T_-values were excluded from the analysis. *: *p* ≤ 0.05, **: *p* ≤ 0.01, ****: *p* ≤ 0.0001.

As expected, male and female Mdr2^-/-^ mice at 26 and 52 weeks of age exhibit a significant up-regulation of *Col1a1* expression in the liver (Figure 2B). Additionally, Mdr2^-/-^ livers displayed a notable increase in *Mcpt2* compared to matching WT controls (Figure 2C). Furthermore, we found a significant correlation between the strength of liver fibrosis (as indicated by *Col1a1* expression) and the expression of MC marker genes *Kit*, *Mcpt1* and *Mcpt2*. In contrast, *Fcer1a*, *Cpa3*, *Tpsb2*, *Mcpt4* and *Mcpt5*/*Cma1* were not significantly co-regulated with liver fibrosis progression in Mdr2^-/-^ mice (Figure 2D).

Altogether, we were able to demonstrate in two independent murine liver fibrosis models that the expression of several MC markers correlates with the severity of disease progression. We further confirmed our experimental findings by analysing public data on an alternative fibrosis mouse model (CCl_4_, see reference (20)). In that study, Holland *et al.* investigated the effects of chronic CCl_4_ treatment on liver fibrosis progression and transcriptome in WT mice. We used data set GSE167216 to examine the expression of MC markers in response to chronic CCl_4_ treatment.

Initial analysis of this data set revealed that the expression of MC markers *Cpa3*, tryptase alpha-1 and tryptase beta-1 (both encoded by *Tpsab1*) and *Tpsb2* correlates with each other in murine liver fibrosis, indicating the presence of MCs in the respective liver tissues (Supplementary Figure 2A). Mice with high expression of MC markers *Fcer1a*, *Cpa3*, *Tpsb2*, *Cma1,* and *Tpsab1* exhibited significantly stronger liver fibrosis compared to the low-expression cohort (Figure 3A). Consequently, we found a significant correlation between these MC markers and *Col1a1* expression (Figure 3B). Notably, *Kit* expression did not correlate with liver fibrosis in this data set (Figure 3A-B).

**Figure 3.**
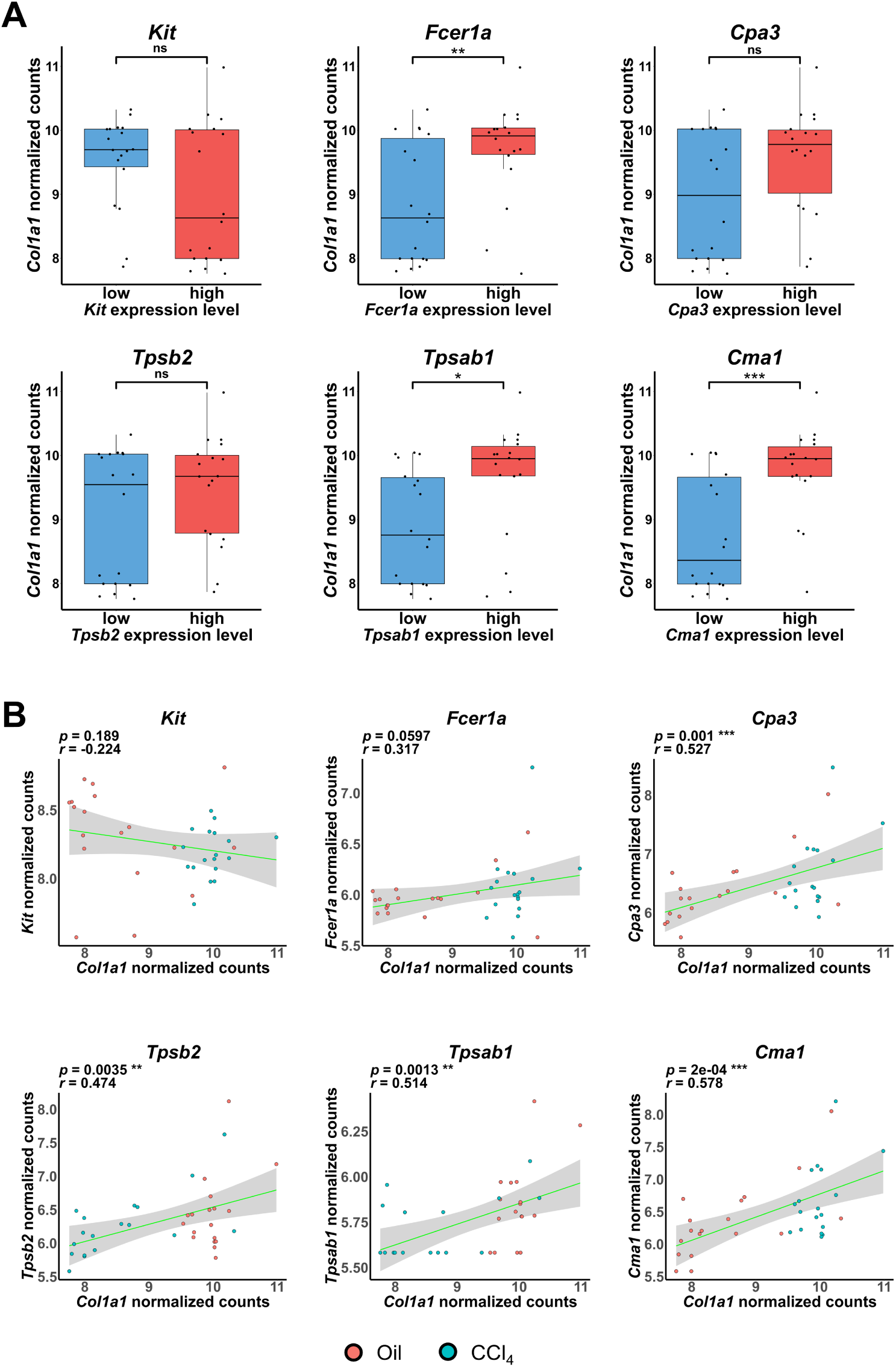
Independent data from CCl_4_–treated mice confirm the up-regulation of MC-specific genes in murine liver fibrosis. After count normalization using variance stabilizing transformation (VST), transcription levels of six MC-specific genes (*Kit*, *Fcer1a*, *Cpa3*, *Tpsb2*, *Tpsab1*, *Cma1*) as well as *Col1a1* were analyzed from the data set GSE167216. This dataset describes the transcriptome of 36 C57BL/6N mice treated with either CCl_4_ or olive oil for 2, 6 or 12 months. (**A**) Mice were categorized into cohorts with high (above median) or low (below median) expression of *Kit*, *Fcer1a*, *Cpa3*, *Tpsb2*, *Tpsab1*, and *Cma1*. The severity of liver fibrosis was compared between both groups using *Col1a1* expression as a key indicator. Data are presented as boxplots with the central line indicating the median, box edges representing the interquartile range (IQR), and whiskers extended to 1.5 time the IQR. Individual points beyond the whiskers are considered as outliers. (**B**) Correlation analysis of *Col1a1* expression with the expression of MC-specific genes (*Kit*, *Fcer1a*, *Cpa3*, *Tpsb2*, *Tpsab1*, *Cma1*). *r*: Pearson correlation coefficient; ns: not significant, *: *p* ≤ 0.05, **: *p* ≤ 0.01, ***: *p* ≤ 0.001.

### 3.3 The correlation between MC marker expression and liver fibrosis progression shows strong similarities between mice and humans

The focus of the present study was to investigate the potential role of MCs in murine liver fibrosis. Additionally, we aimed to determine if the correlations found between MC-specific gene expression and fibrosis progression in mice could be applied to humans. Since liver fibrosis is considered a precursor to hepatocarcinogenesis (26), we analyzed data from patients with liver cancer from The Cancer Genome Atlas Program (TCGA) (https://portal.gdc.cancer.gov/projects/TCGA-LIHC) similar to the data presented in Figure 3. Consistently, we confirmed a correlation of *Cpa3*, *Tpsab1,* and *Tpsb2* expression in human HCC tissue (Supplementary Figure 2B). Furthermore, we observed a strong correlation between fibrosis progression and the expression of human MC marker genes *KIT*, *FCER1A*, *CPA3*, *TPSB2* and *TPSAB1* (Figure 4A-B). Notably, *CMA1*, the human equivalent of the murine *Mcpt5*/*Cma1* gene, did not show co-regulation with liver fibrosis progression in patients, as was found for its murine counterpart in two out of three mouse models studied (compare Figure 1D, 2D).

**Figure 4.**
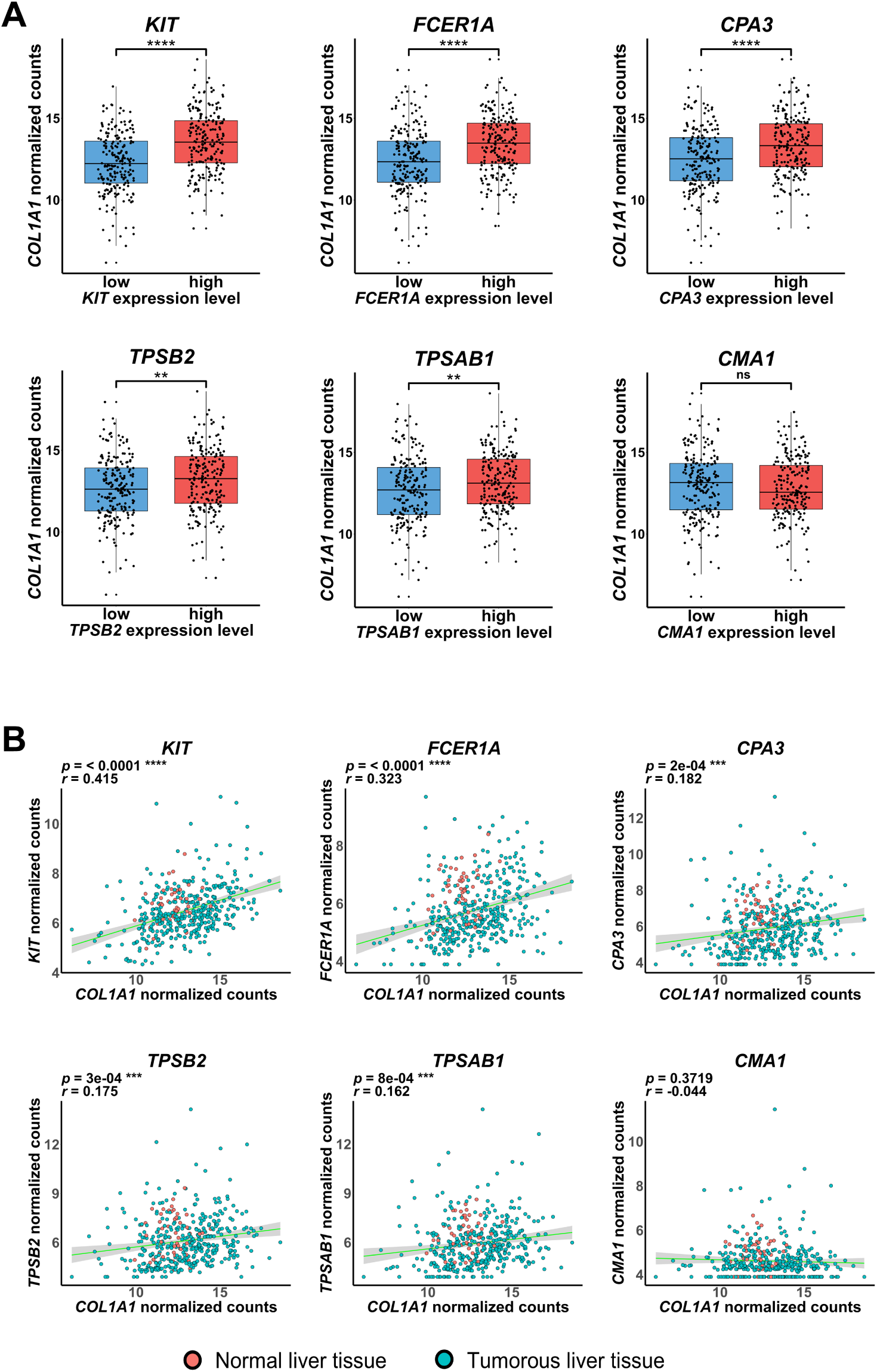
The correlation between MC marker expression and liver fibrosis progression shows strong similarities between mice and humans. Bioinformatic analysis was conducted on 371 human HCC samples and 50 matching samples of surrounding non-tumorous liver tissue from The Liver Hepatocellular Carcinoma (LIHC) dataset in The Cancer Genome Atlas (TCGA). Patient-derived mRNAseq data was examined for *COL1A1* expression, as well as the expression of MC markers *KIT*, *FCER1A*, *CPA3*, *TPSB2*, *TPSAB1* and *CMA1*. (**A**) Patients were divided into groups with high (above median) or low (below median) expression of MC markers, similar to the analysis in Figure 3A. The level of liver fibrosis in both groups was then assessed by *COL1A1* expression and displayed as boxplots. (**B**) Correlation analysis was performed on *COL1A1* expression and the expression of selected MC-specific genes in HCC samples (green) and normal liver tissue (red). *r*: Pearson correlation coefficient; ns: not significant, ** : *p* ≤ 0.01, *** : p ≤ 0.001; **** : *p* ≤ 0.0001.

We expanded our analysis of patient data from the TCGA-LIHC data set and examined the differential gene expression between patients with a high or low score of an MC expression signature (*TPSB2*, *CPA3* and *TPSAB1*). Interestingly, these patients also exhibited significantly increased expression of other MC-relevant genes not included in the signature, as well as genes generally associated with fibrogenesis and inflammation (Supplementary Figure 3A). A selection of enriched gene ontology terms linked to significantly up-regulated genes in these patients is displayed in Supplementary Figure 3B. Furthermore, genes linked to these enriched terms and their intersection across various up-regulated biological processes are illustrated in Supplementary Figure 3C.

### 3.4 Hepatic MCs from mice can be purified by FACS and typically do not express Mcpt5*/*Cma1

To utilize mouse models for studying MC biology in the liver and their role during liver fibrosis, selective genetic manipulation in MCs would be desirable. Sasaki *et al.* have recently generated mice in which the human diphtheria toxin A (DT) receptor is expressed under the control of the endogenous *Mcpt5* promoter. The authors claim almost complete depletion of MCs in these mice upon DT injection (27). These data suggest that the *Mcpt5* promoter might be suitable for transgenic control of MC-specific gene expression. *Mcpt5*-*Cre* transgenic mice have already been described and are able to drive MC-specific reporter gene expression at least in murine peritoneal cavity and skin (16). However, our data from the present study show that, in contrast to other MC-markers tested, *Mcpt5* expression levels in the mouse liver do not correlate well with fibrosis severity. Thus, the impact of *Mcpt5* in the mouse liver is unclear.

Therefore, in this study, we aimed to determine if *Mcpt5*-*Cre* transgenic mice are suitable for gene targeting in hepatic MCs. To do this, we first crossed the *Mcpt5*-*Cre* transgene into mT/mG reporter mice (17), resulting in animals referred to as mT/mG; *Mcpt5-Cre* mice. The concept of these reporter mice is shown in Figure 5A. Additionally, we crossed these mice with Mdr2^-/-^ mice (Mdr2^-/-^ mT/mG; *Mcpt5-Cre*) to conduct MC-specific lineage tracing in fibrotic livers.

**Figure 5.**
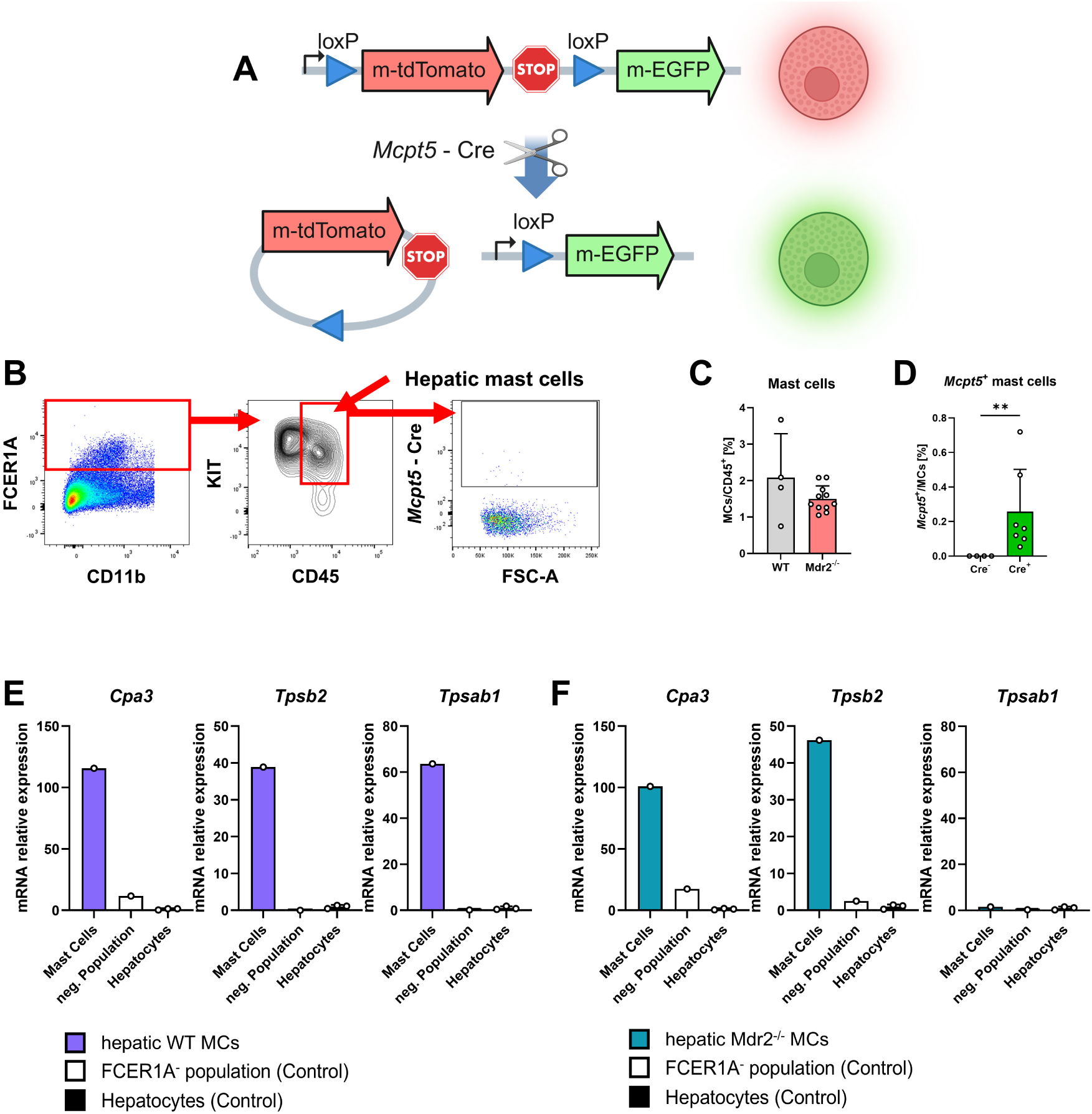
Establishment of a FACS-based protocol for isolating MCs from murine liver and their transcriptional characterization by qPCR. (**A**) Experimental setup and mouse strains. We used Mdr2^-/-^ mice, which develop spontaneous hepatic fibrosis and their wildtype (WT) counterparts. To visualize *Mcpt5*/*Cma1* expressing MCs, we additionally employed Mdr2^-/-^mT/mG; *Mcpt5*-*Cre* mice. In these reporter mice, cells lacking *Mcpt5* expression show red fluorescence (tdTomato), whereas *Cre*-mediated excision of the floxed tdTomato cassette in *Mcpt5*^+^ cells leads to expression of EGFP and thus green fluorescence. (**B**) Gating strategy for MC characterization by flow cytometry. After removing doublets and gating on immune cells (CD45^+^), liver MCs were identified as FCER1A/KIT double-positive and CD45^high^ cells. Subsequent gating on EGFP-positive cells enables the assessment of *Mcpt5*-expression. (**C**) Efficacy of MC isolation. The percentage of MCs relative to total CD45-positive leucocytes is shown for WT and *Mdr2*^-/-^ mT/mG mice. Each data point represents one liver. Please note that the *Mdr2*^-/-^ mT/mG group contains both *Mcpt5*-*Cre*-positive (n = 7) and negative (n = 4) animals. (**D**) Quantification of *Mcpt5*-expressing hepatic MCs in fibrotic livers. Analyzed MCs from (C) (n = 11) were categorized into *Mcpt5-Cre*-positive and -negative groups and examined for EGFP expression. **: *p* ≤ 0.01. Percentages reflect the fraction of *Mcpt5*-expressing MCs within total isolated MCs. (**E**-**F**) Transcriptional characterization of FACS sorted MCs by qPCR. Hepatic MCs were isolated by FACS sorting (Gating strategy shown in panel B) and subjected to mRNA isolation and qPCR for MC-markers *Cpa3*, *Tpsb2* and *Tpsab1.* FCER1A-negative cells and hepatocytes served as negative controls. (**E**) Results for hepatic MCs from 52-week-old WT mice. (**F**) Hepatic MCs from 12-week-old Mdr2^-/-^ mice. Expression values were normalized to *Gapdh* and presented as fold induction relative to the basal expression in hepatocytes.

As a proof-of-concept pilot experiment, we cultured bone marrow-derived mast cells (BMMCs) from mT/mG; Mcpt5-Cre and WT mice *in vitro* following a protocol that typically yields >95% KIT (CD117) and FCER1A double-positive MCs (19). Using this approach, we were able to identify a subset of EGFP-positive reporter BMMCs through immunofluorescence microscopy (Supplementary Figure 4A) and FACS analysis (Supplementary Figure 4B), demonstrating the functionality of the reporter system.

We then established an isolation protocol and a FACS-based gating strategy for the detection and purification of primary MCs from mouse liver (Supplementary Figure 5). Using this protocol, we isolated hepatic MCs from WT and Mdr2^-/-^ mT/mG; *Mcpt5*-*Cre* mice. In WT liver, we found that approximately 2% of analysed CD45^+^ leukocytes are KIT^+^/FCER1A^+^ MCs (Figure 5B-C). Similar proportions of MCs were found in Mdr2^-/-^ mT/mG mice with liver fibrosis that were either *Mcpt5*-*Cre* positive (n=7) or *Cre*-negative (n=4, Figure 5C). However, on average, only about 0.2% of MCs from Mdr2^-/-^ mT/mG; *Mcpt5*-*Cre* liver were EGFP-positive and thus expressed *Mcpt5* (Figure 5D). From this data we conclude that *Mcpt5* is not expressed in murine hepatic MCs.

Our next goal was to purify MCs on a larger scale to allow subsequent molecular analysis, *e.g.* by qPCR. We thus isolated liver immune cells from either WT or Mdr2^-/-^ mice and enriched KIT^+^/FCER1A^+^ MCs using a FACSAria™ Fusion cell sorter (BD Biosciences). This approach enabled the isolation of MCs in the range of 70,000 – 150,000 cells per mouse liver. By pooling these cells (n= 3-5), we were able to isolate sufficient RNA to demonstrate by qPCR that hepatic murine MCs do express typical MC marker genes such as *Cpa3* and *Tpsb2* in both WT and Mdr2^-/-^ livers, while *Tbsab1* was only expressed in hepatic WT MCs, but not in MCs derived from Mdr2^-/-^ liver (Figure 5E-F).

### 3.5 Identification of hepatic murine MCs *in situ* by Molecular Cartography™

Our previous results strongly suggested that in the murine liver, MCs do exist and increase in number during chronic liver fibrosis and HCC. However, the *in situ* confirmation of MCs in the injured mouse liver using conventional staining approaches had not been convincingly successful. To address this issue, we applied a spatial transcriptomics approach known as Molecular Cartography™ (Resolve BioSciences GmbH, Monheim, Germany), as recently described (24). In this analysis, we examined liver tissue sections from Mdr2^-/-^ mice and identified MC-specific transcripts. Cells showing evidence of transcripts specific for at least two different MC-specific genes were classified as MCs. Through this method, we successfully identified hepatic murine MCs *in situ* in Mdr2^-/-^ tissue at an average frequency of 2 cells/mm^2^ (Figure 6A-C).

**Figure 6.**
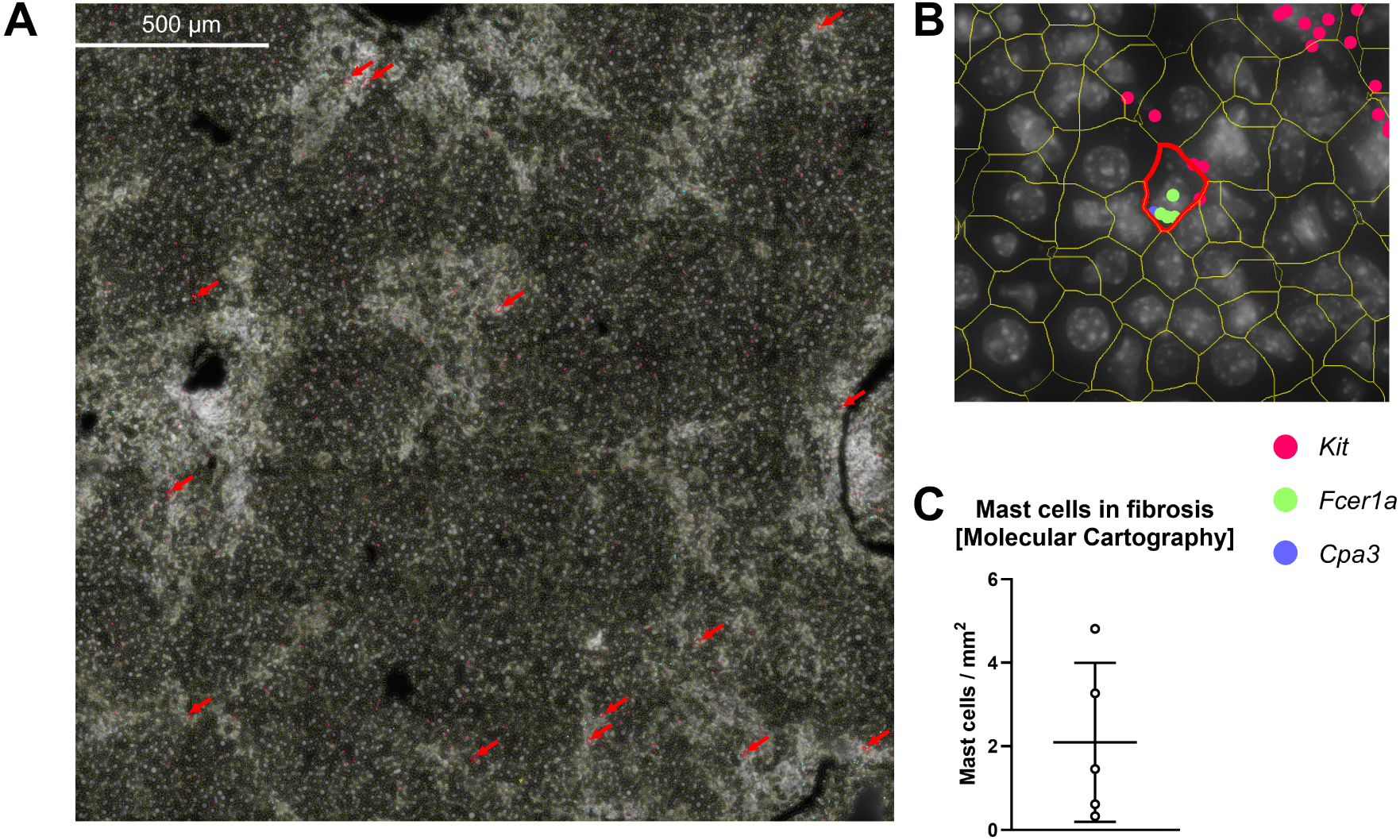
Use of Molecular Cartography™ to identify mast cells in the murine liver. Liver FFPE sections from WT mice (n=1) or Mdr2^-/-^*Casp8*^Δhepa^ mice (n=4), one mouse with two sections) were analyzed using Molecular Cartography™. (**A**) Histologic overview of a representative DAPI-stained tissue section from an Mdr2^-/-^ liver. DAPI images were used to define cell segmentation overlay followed by cell-wise transcript identification. Cells expressing *Kit* or *Cpa3,* along with at least one other MC-specific gene such as *Fcer1a* were identified as MCs and are highlighted by red arrows. (**B**) Enlarged view of a representative liver tissue section from (A), showing a MC with mRNA co-expression of *Kit*, *Fcer1a,* and *Cpa3* highlighted by red boundaries. (**C**) Quantification of total MCs from all (n = 5) analyzed fibrotic liver sections per mm^2^ in our study. Data are presented as mean ± SD.

Overall, this data provides new evidence that MCs do exist in the murine liver, albeit at a very low frequency, and can be visualized using spatial transcriptomics or FACS after tissue dissociation. This established methodology will enable the precise characterization of hepatic MC populations in mice, such as through single-cell RNA sequencing approaches, and support further in-depth analysis of MC biology in murine liver fibrosis models.

## 4 Discussion

Mast cells (MCs) are heterogeneous myeloid-derived immune cells known for their pro-inflammatory role in allergies (28). More recently, it has become evident that MCs may also play important roles in the progression of liver disease, including liver fibrogenesis and hepatocarcinogenesis, as reviewed by us and others (2, 29). However, most available data on this topic was collected from patients and sometimes from rats, while studies on mice are rather rare. Therefore, basic scientific studies on the function of MCs in the mouse liver would be particularly valuable, especially if combined with the powerful possibilities of conditional, MC-specific gene targeting (*e.g.* cre/loxP system). Challenges with the use of mouse models in MC research include the presumed low absolute number of resident MCs in the liver. Additionally, some key studies on hepatic MCs in mice have been conducted in the Mdr2^-/-^model of cholestatic liver fibrosis and inflammation-associated HCC (14), which have recently been retracted (30, 31). Thus, the current state of knowledge on the biology of hepatic MCs in mice is low and needs to be critically revised. In particular, the following questions were newly posed and mostly answered in the course of the present study:

i. Are there any resident mast cells in the mouse liver?
ii. Do MCs play a role in the progression from chronic liver damage to liver fibrosis?
iii. Do hepatic MCs in mice have a tissue-specific gene signature?
iv. How can the assumed small number of murine hepatic MCs be investigated systematically?

Throughout the present study, we were unable to detect hepatic MCs in mice with liver fibrosis using conventional MC-staining protocols such as Toluidine blue staining, which has been approved in murine skin (6, 32), or immunofluorescence staining with a combination of MC-specific antibodies as demonstrated in murine dural meninges (33). Only through spatial transcriptomics approaches were we able to visualise MCs by detecting intracellular, MC-specific transcripts in the mouse liver. Using this method, we estimated an average frequency of 2 MCs/mm^2^ in an adult, fibrotic Mdr2^-/-^ mouse liver. The reason why we were unable to stain hepatic mast cells *in situ* using established dyes or MC-specific antibodies remains unclear, particularly since the methods used gave satisfactory results in different tissues. It is possible that during the removal and processing of mouse liver, most of the MC-proteins may be destroyed by unknown mechanisms, while MC mRNA in liver tissues appears to be more stable and detectable Another limitation was the lack of available mouse- and MC-specific antibodies validated for tissue staining. This could be a crucial issue, as we were able to detect hepatic MCs using KIT and FCER1A antibodies in isolated MCs from mouse liver in FACS analyses. Therefore, it may be worthwhile to invest more effort in developing improved staining methods for visualizing hepatic MCs in the future.

Initially, the direct detection and quantification of MCs in the fibrotic mouse liver was not possible. Therefore, we developed indirect methods to investigate the role of MCs in the progression of liver fibrosis in mice. To achieve this, we utilized murine liver samples from two well established HCC-associated liver fibrosis models: DEN/CCl_4_ treatment (generating toxic chronic injury plus pro-carcinogenic stimulation) and genetic Mdr2 deletion (Mdr2^-/-^ mice, develop chronic cholestasis). Additionally, we analysed publicly available liver gene expression data from a pure CCl_4_ liver fibrosis model published by Holland and colleagues (20). Overall, we observed a clear and significant correlation between the expression of several MC markers and the progression of liver fibrosis, albeit with some model-specific differences. These correlations are schematically summarized in Figure 7.

**Figure 7.**
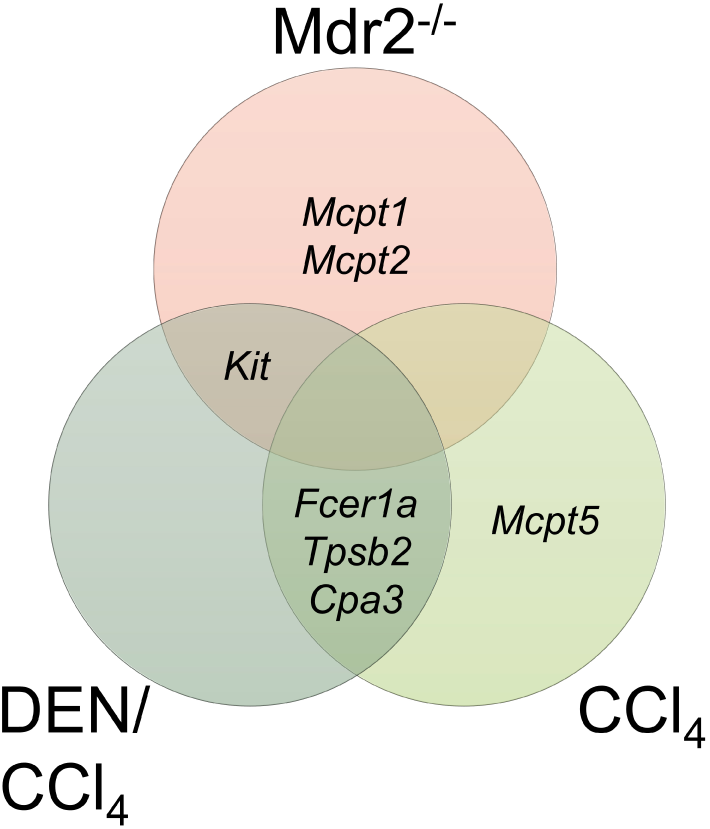
Venn Diagram related to the data shown in Figure 1-3. The expression of MC marker genes was correlated with the expression of the *Col1a1* gene in three different mouse models of liver fibrosis: Mdr2^-/-^ (genetic deletion of the Mdr2 gene), DEN/CCl_4_ (chemical induction of liver fibrosis and HCC due to treatment with diethylnitrosamine and carbon tetrachloride), and CCl_4_ (liver fibrosis induction through CCl_4_ treatment only). Only MC markers that significantly correlate with the expression of *Col1a1* in one or more models are shown. Please note that this diagram does not display the expression levels of the MC markers themselves. Therefore, the absence of e.g. *Kit* or *Fcer1a* in a distinct section does not imply that *Kit* / *Fcer1a* are not expressed in these mice. Instead, *Kit* / *Fcer1a* expression in the respective cohorts does not correlate with fibrosis severity.

It was found that none of the tested MC marker expressions correlated in all three injury models with the level of fibrosis progression (measured by *Col1a1* gene expression). This indicates that the MC signature in the fibrotic mouse liver is dependent on the type of the damaging noxious agent. This seems plausible, as the general tissue environment and the microenvironment in CCl_4_-treated livers (toxic injury) (34) is very different compared to the tissue environment in Mdr2^-/-^ mice (cholestatic injury) (35).

However, when only the CCl_4_-associated, toxic mouse models (DEN/CCl_4_ *versus* CCl_4_) are considered, a common group of three MC-related genes *Fcer1a*, *Tpsb2 and Cpa3* can be defined, whose expression levels correlate significantly with *Col1a1* gene expression. Additionally, CCl_4_-induced murine liver fibrosis progression correlates with *Mcpt5/Cma1* expression (encoding mast cell chymase 1). In sharp contrast, liver fibrosis progression in Mdr2^-/-^ mice was associated with increased expression of *Kit*, *Mcpt1* and *Mcpt2*, but remarkably not with *Mcpt5*. In good agreement, our further investigations at the single-cell level showed that MCs isolated from wildtype or Mdr2^-/-^ mouse liver generally do not express *Mcpt5/Cma1*. Additionally, primary MCs from Mdr2^-/-^ liver express *Tpsb2* (encoding tryptase beta-2) and in WT liver *Tpsab1* (encoding tryptase alpha-1 and beta-1).

Classically, two subtypes of MCs are distinguished in mice and humans based on their localisation, function and gene expression signature: mucosal mast cells (MMCs) and connective tissue mast cells (CTMCs) (36). However, very recent results based on single-cell RNA sequencing (scRNAseq) by Tauber and colleagues (8) show that the diversity of MC subpopulations is significantly higher. The consensus is that MMCs express *Mcpt1* and *Mcpt2*, but not *Mcpt4* or *Mcpt5*. *Cpa3* expression, which correlated with the degree of liver fibrosis in toxic (*i.e.* CCl_4_, DEN/CCl_4_) injury models investigated, is obviously expressed in all murine MC subtypes, as transgenic expression of Cre recombinase in the *Cpa3* locus (*i.e. Cpa3*^+/Cre^ mice) is able to deplete all mouse MCs, although it was shown that *Cpa3* as well as *Tpsb2* is more strongly expressed in CTMCs (37).

In conclusion, our findings suggest that MCs in fibrotic liver of Mdr2^-/-^ mice exhibit some characteristics of mucosal MCs, whereas MCs from CCl_4_-treated mice are more similar to connective tissue MCs. Future in depth analyses (*e.g.* scRNAseq) of purified MCs from fibrotic as well as tumour-bearing liver samples will be necessary to more thoroughly depict the diversity of liver MCs. Such analyses will now be feasible as we have developed the methodological tools to isolate the necessary number of MCs from mouse liver in the present study.

Our correlation analyses and proposed MC signatures in HCC-associated mouse fibrosis models align well with similar analyses in the TCGA HCC patient cohort (https://portal.gdc.cancer.gov/projects/TCGA-LIHC). Once again, no correlation between fibrosis progression and *CMA1*/Mcpt5 expression could be identified, suggesting that MCs in the human fibrotic liver share similarities with murine hepatic MCs. Additionally, we demonstrated in the human HCC data that high expression of an MC gene signature was linked to the up-regulation of multiple genes associated with fibrosis and inflammatory pathways. However, it is important to note that this patient population is highly diverse, with various disease aetiologies, making comparisons with the standardised mouse models challenging.

We successfully detected MCs in the mouse liver using spatial transcriptomics and purified them with optimised cell isolation protocols and FACS methods. This technical advancement will serve as a prerequisite for future in-depth analysis of hepatic MCs in mice. Through extensive correlation analyses from three liver injury models, we have shown that the expression levels of established MC markers are significantly correlated with the degree of liver fibrosis. Therefore, these MC markers have the potential to be used as additional biomarkers for assessing liver fibrosis. The expression signature of these markers suggests that murine liver MCs are heterogeneous but share many properties with mucosal mast cells. Overall, targeting MCs in the liver could be a complementary approach to treating liver fibrosis in the future.

## 5 Acknowledgements

This work was supported by the German Research Foundation (DFG), grant LI1045/6-1/WE2554/15-1/HU794/ME3431 (Project No. 452602471) awarded to C.L., R.W., M.H. and S.K.M. C.L. was further supported by DFG grants LI1045/4-2 (Project No. 271777553) and DFG/CRC 1382 (Project A02, Project No. 403224013). Additionally, this work received support from the Flow Cytometry Facility of the Interdisciplinary Center for Clinical Research (IZKF) within the Faculty of Medicine at RWTH Aachen University and projects funded by the IZKF (grants PTD1-5 and PTD1-6).

## 6 Author contributions

C.P. developed and performed MC isolation protocols, along with conducting bioinformatic, statistical, and Molecular Cartography™ data analysis. J.O. conducted animal experiments. C.L., R.W., M.H. and S.K.M. secured funding, designed experiments, and carried out data analysis and interpretation. C.L. and C.P. drafted the manuscript and supervised the study, with all authors reading, approving, and editing the final version of the paper.

## 7 Ethical statement

All animal experiments were conducted in compliance with German legal requirements and approved by the authority for environment conservation and consumer protection of the state of North Rhine-Westphalia (LANUV).

## Supplementary figures

**Supplementary Figure 1.**
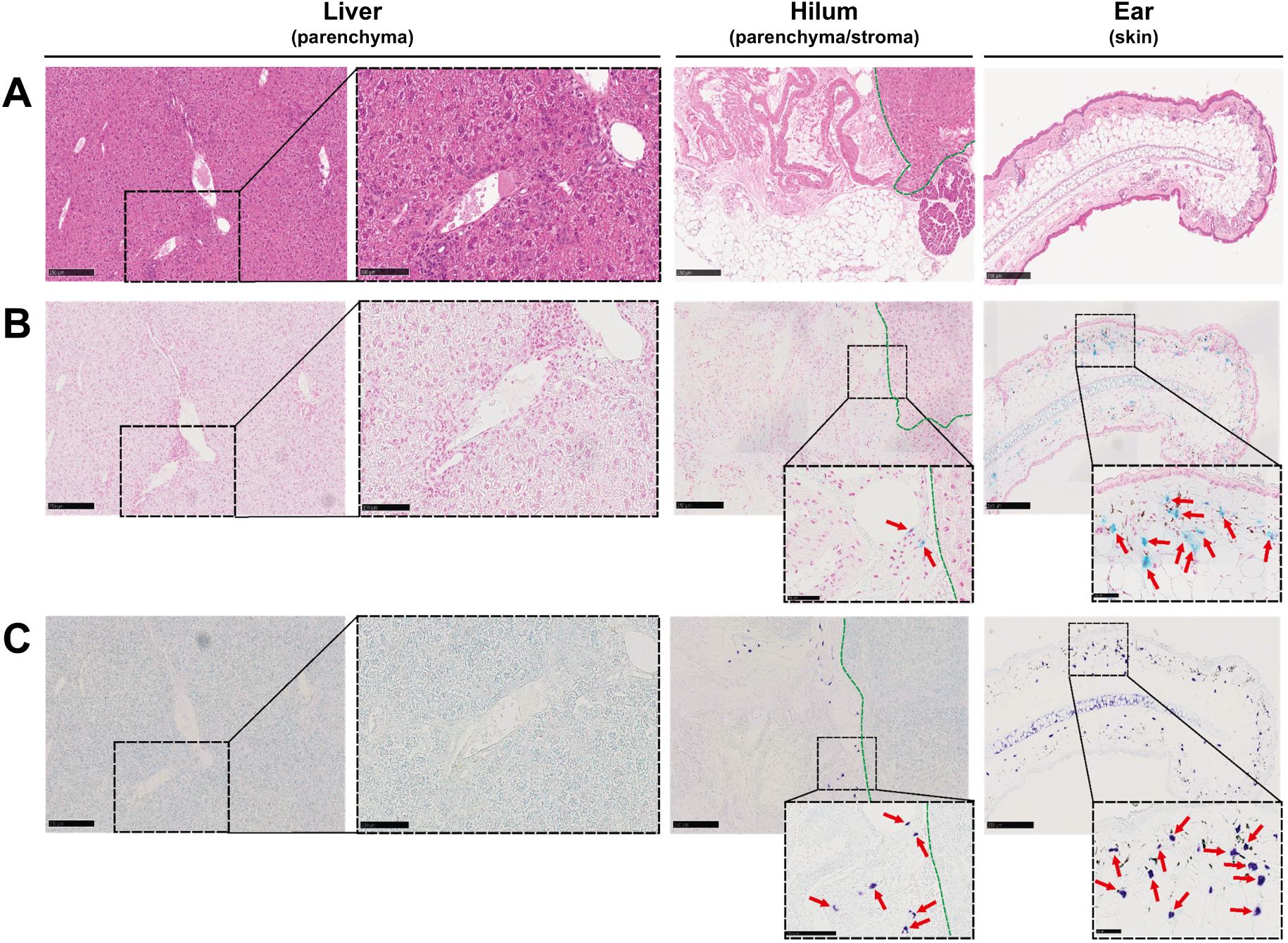
MC-specific immunohistological staining methods established on human tissue samples are not applicable in murine liver. (**A-C**) Formalin fixed, paraffin embedded serial tissue sections of fibrotic murine liver (Mdr2^-/-^), the hilum area including parenchyma and stromal tissue (separated by a green dashed line) and skin as control tissue (mouse ear) were stained by (**A**) hematoxylin and eosin, (**B**) Alcian blue or (**C**) Toluidine blue to visualize mast cells. Stain-positive mast cells were exclusively found in control tissues (stroma and skin) and are highlighted by red arrows in enlarged image details.

**Supplementary Figure 2.**
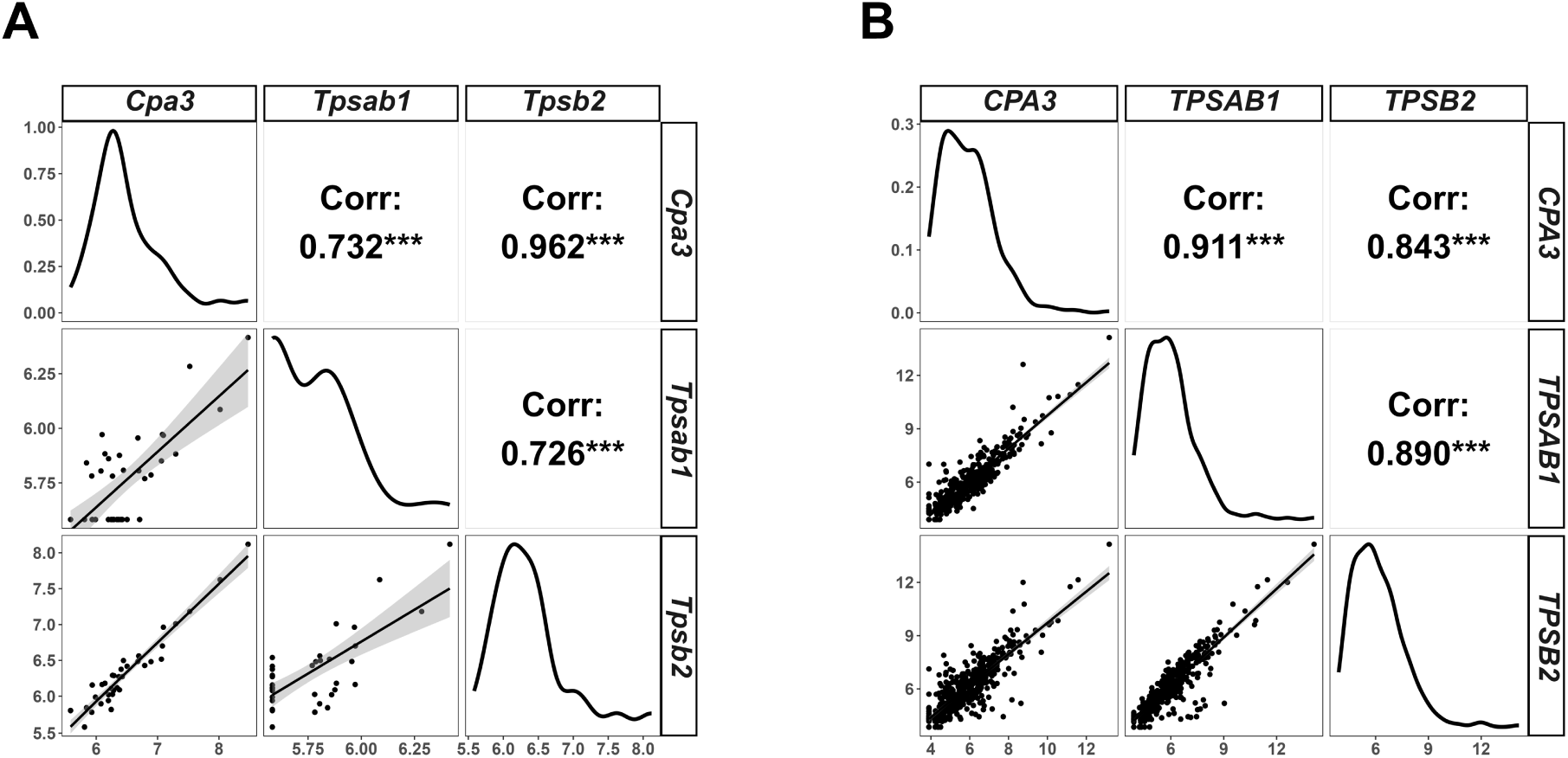
Expression of MC markers *Cpa3*, *Tpsab1* and *Tpsb2* correlate with each other in murine liver fibrosis and in liver samples from HCC patients. (**A**) Figure related to Figure 3. VST normalized gene expression data for *Cpa3*, *Tpsab1* and *Tpsb2* in murine CCl_4_-induced liver fibrosis was extracted from the data set GSE167216. The Pearson correlation matrix for *Cpa3*, *Tpsab1* and *Tpsb2* is shown. The distribution of expression values is displayed for each gene on the diagonal. In scatter plots below the diagonal, individual measurements of gene expression are plotted against each other, with a regression line and confidence interval (gray area). Pearson correlation coefficients with significance levels are provided in the fields above the diagonal. (**B**) Figure related to Figure 4. VST normalized gene expression data for *CPA3*, *TPSAB1* and *TPSB2* in human HCC samples was extracted from the TCGA-LIHC dataset and displayed as Pearson correlation matrix as in (A). *** : p ≤ 0.001.

**Supplementary Figure 3.**
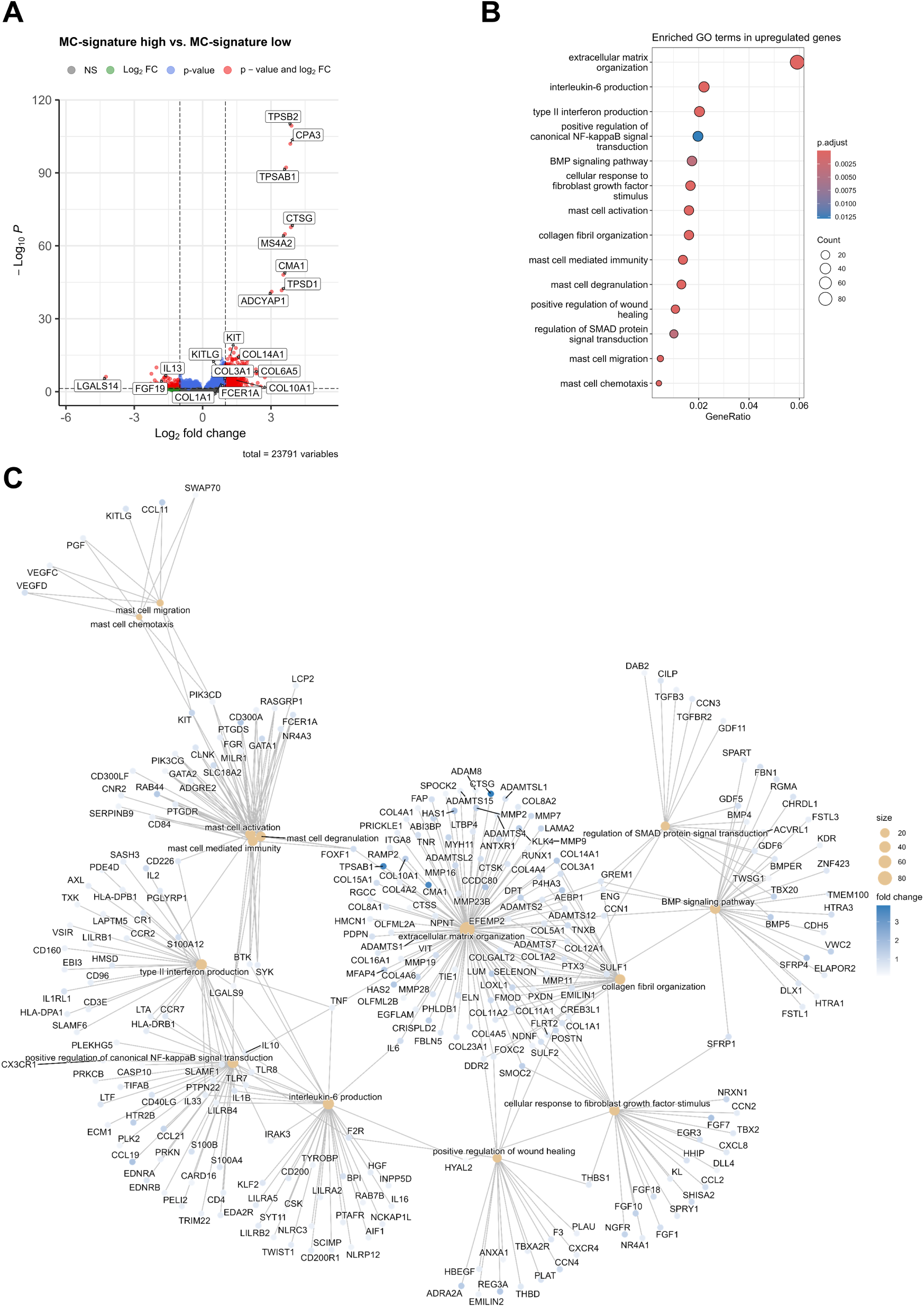
High mast cell signature expression in HCC is associated with increased expression of inflammation- and fibrosis-associated genes. (**A**) Volcano plot showing differential up- and down-regulated genes between HCC samples with high or low MC gene signature expression. Log_2_ -fold changes are given on the x-axis while −log_10_ *p*-values (adjusted *p*-values) are shown on the y-axis. The horizontal dashed line indicates the significance level *p* = 0.05, and vertical dashed lines indicate a 2-fold change of gene expression (log_2_(1)). Selected genes are annotated by arrows. (**B**) Dot plot showing selected enriched gene ontology terms (GO) in up-regulated genes in patients with high expression of MC gene signature. (**C**) Network plot of enriched GO terms shown in (B) highlighting upregulated genes and their intersection between different gene ontology terms.

**Supplementary Figure 4.**
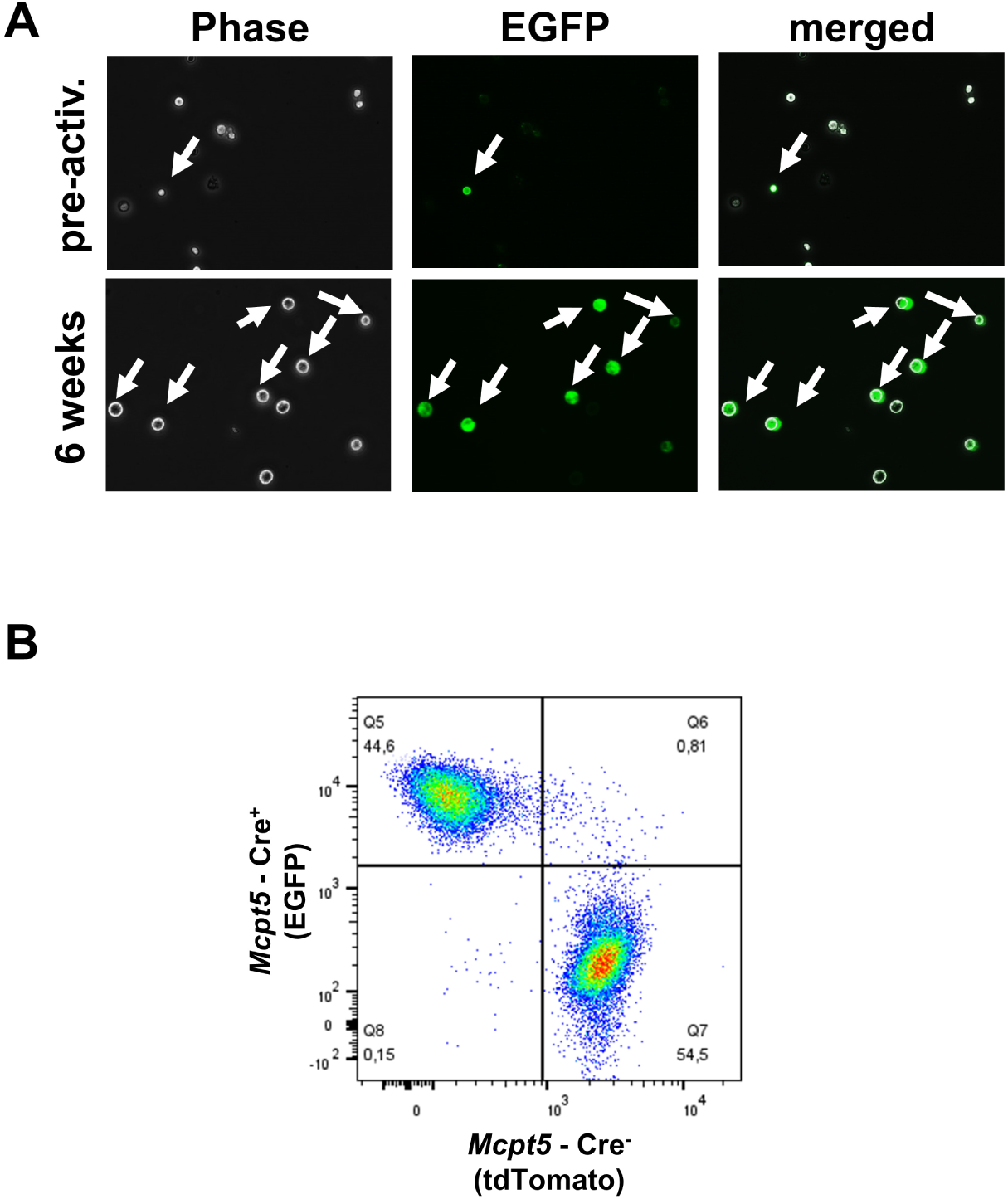
Proof of functionality of mT/mG; *Mcpt5*-*Cre* reporter mice. (**A**) BMMCs were generated from mT/mG; *Mcpt5*-*Cre* reporter mice (compare Figure 5A) and imaged before (pre-activ.) and after six weeks of activation with IL-3 to assess EGFP expression. (**B**) BMMCs generated from mT/mG; *Mcpt5*-*Cre* reporter mice were indirectly analyzed for their expression of *Mcpt5*-Cre by FACS. CD45^+^ single cells were characterized as Cre^+^ (EGFP^+^ tdTomato^-^) or Cre^-^ (EGFP^-^ tdTomato^+^).

**Supplementary Figure 5.**
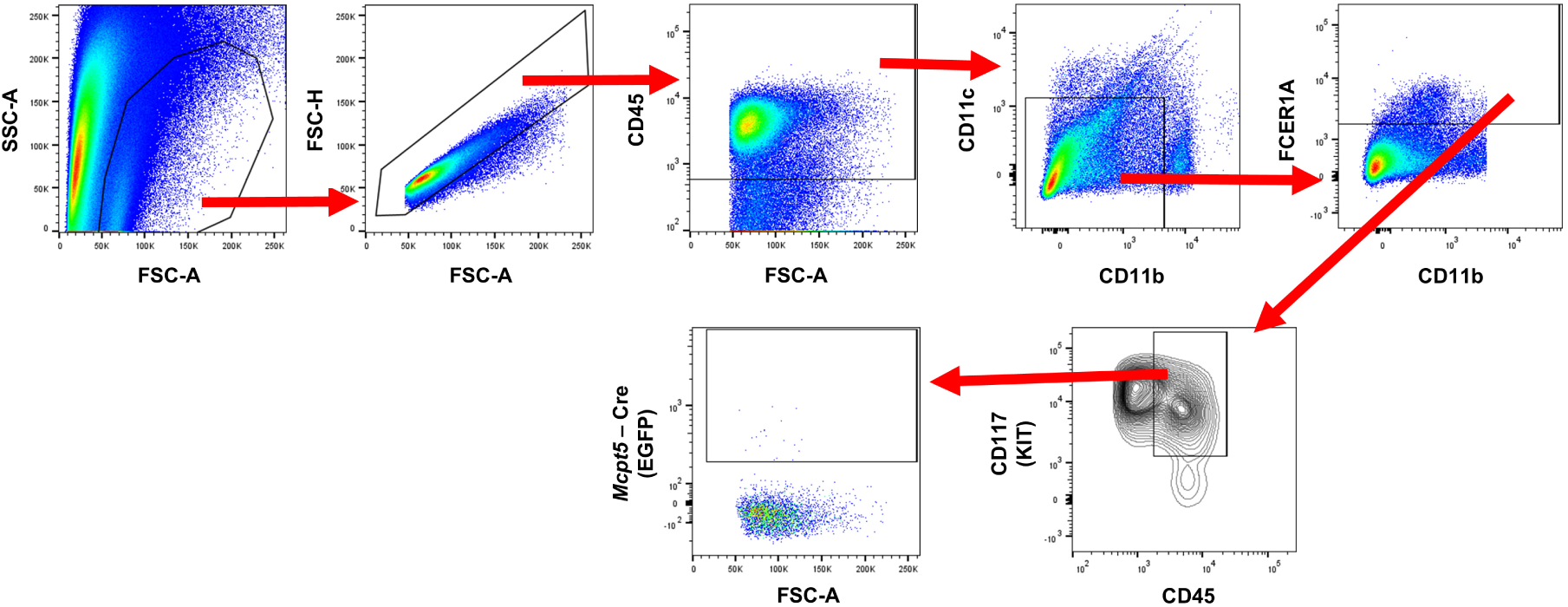
FACS gating strategy to detect and purify primary MCs from murine liver. Liver cells were initially selected based on size and granularity (FSC-A vs. SSC-A) and then doublets were removed (FSC-A vs. FSC-H). The immune cell population (CD45^+^) was further narrowed down to CD11c^-^ and CD11b^low-intermediate^ cells and MCs were gated as FCER1A^+^ and CD117^+^ CD45^high^. MCs isolated from *Mcpt5 -* reporter mice were additionally characterized by their expression of *Mcpt5* - Cre as determined in the EGFP channel.

